# Neural and behavioral reinstatement jointly reflect retrieval of narrative events

**DOI:** 10.1101/2024.10.19.619187

**Authors:** Matthias Nau, Austin Greene, Hannah Tarder-Stoll, Juan Antonio Lossio-Ventura, Francisco Pereira, Janice Chen, Christopher Baldassano, Chris I. Baker

## Abstract

When recalling past events, patterns of gaze position and neural activity resemble those observed during the original experience. We hypothesized that these two phenomena, known as gaze reinstatement and neural reactivation, are linked through a common process that underlies the reinstatement of past experiences during memory retrieval. Here, we tested this proposal based on the viewing and recall of a narrative movie, which we assessed through fMRI, deep learningbased gaze prediction, and language modeling of spoken recall. In line with key predictions, gaze behavior adhered to the same principles as neural activity; it was event-specific, robust across individuals, and generalized across viewing and recall. Additionally, gaze-dependent brain activity overlapped substantially across tasks. Collectively, these results suggest that retrieval engages mechanisms that direct our eyes during natural vision, reflecting common constraints within the functional organization of the nervous system. Moreover, they highlight the importance of considering behavioral and neural reinstatement together in our understanding of remembering.

## Introduction

Eye movements determine the content, spatial organization, and relative timing of all visual impressions we obtain of our environment. Simultaneously, visual inferences guide our eyes towards behaviorally relevant cues (e.g., when recognizing a face through sequential sampling). Gaze behavior is therefore a fundamental component of vision (1–3), with viewing statistics necessarily shaping the activity of all visually responsive circuits in the brain (e.g., gaze shifts drive activity fluctuations (4)). Even structures typically associated with memory, such as the hippocampus, show retinotopic activity modulations (5–9) as well as eye-movement signals (10–14), implying that the connection to gaze behavior extends beyond putative boundaries of what is commonly referred to as the visual system.

Importantly, the way neural circuits engage during active vision likely also constrains their involvement in other tasks, such as recall. This is because a circuit’s activity patterns are inextricably tied to its anatomy, which reflects the circuit’s engagement over developmental and evolutionary timescales (15, 16). Because this engagement is inherently linked to gaze behavior, the functional organization of widespread neural circuits likely embodies how we move through and sample the environment during natural vision. As a consequence, many principles invoked for active vision should generalize to memory retrieval and episodic simulation (i.e., phenomenological experiences of past, future, or fictitious events in the absence of physical stimuli (17, 18)). These general principles should include sequentiality, meaning that items are sampled or retrieved in a sequence, with recalled events unfolding over time, as well as the involvement of eye movement-related mechanisms. Preliminary evidence for the existence of such general principles can be found both on the level of behavior and neural activity.

On the behavioral level, eye movements (19–28) and pupil size (29–31) have been shown to reflect imagery, recall, and recognition when probed with simple, static stimuli. In particular, gaze reinstatement describes the observation that gaze patterns during image viewing tend to be recapitulated during retrieval (e.g., for review see (14)). These eye-movement recapitulations have been proposed to play a functional role in retrieval, especially because interfering with them impairs recall and episodic simulation (e.g., (24, 32, 33)). However, to date, it remains unclear whether and how gaze reinstatement extends to the recall of more complex and continuous experiences spanning longer time scales such as those we make in everyday life. Likewise, on the neural level, activity patterns observed during retrieval often resemble those observed during viewing, a phenomenon termed neural reactivation (also referred to as neural reinstatement, for review see e.g., (18, 34, 35)). Using rich and dynamic stimuli, such as movies, neural reactivations have been shown to be specific to individual events that are recalled, and to be consistent across participants recalling the same event (see e.g., (36–41)).

While recent years have seen growing interest in understanding the links between gaze reinstatement and neural reactivation (see e.g., (21, 29, 42–44)), the relationship between these two phenomena remains largely elusive. Here, we hypothesize that gaze reinstatement and neural reactivation are linked through a common process that underlies the reinstatement of past experiences during memory retrieval. Building on ideas and approaches from work on active vision, we test this hypothesis by directly linking patterns in gaze behavior, brain activity, and spoken recall.

## Results

To probe the relationship between gaze reinstatement and neural reactivation in a naturalistic setting, we incorporated camera-based and magnetic resonance-based eye tracking into the “Sherlock Movie Watching Dataset” (36) – the basis of an extensive literature on neural reactivations and their role in recall (e.g., (36, 37, 39, 45–48)). In these data, human volunteers viewed and then recalled a movie while audio recordings captured spoken recall, and while brain activity was monitored with functional magnetic resonance imaging (fMRI). By integrating previously unreleased in-scanner eyetracking data with magnetic resonance-based eye tracking (49) and newly acquired camera-based data from a desktop setup, we were able to test multiple behavior-informed predictions that were previously out of reach.

First, if gaze reinstatement and neural reactivation are indeed linked through a common process, we expect that the two phenomena should not only co-occur in these data, but also share key properties such as event-specificity and robustness across participants (*Prediction 1*). Second, patterns in gaze and neural activity should generalize across viewing and recall within the same dataset (*Prediction 2*). Third, eye movements should correlate with brain activity during viewing and recall, with gaze-dependent activity overlapping between the two tasks (*Prediction 3*).

In the following, we present our study design and the results of testing these predictions in five steps. 1) We start by describing the movie-viewing and recall task as well as our empirical measures of gaze behavior, neural activity, and spoken recall. 2) We next ensure that both datasets are suited for event-specific analyses by showing that participants’ spoken recall followed the event structure of the movie (i.e., the content and order of narrative events), as indicated by language modeling. 3) In line with our first prediction, we then show that gaze patterns during movie viewing are indeed event-specific and highly consistent across participants, which is largely explained by the visual content of the movie. 4) Using a Hidden Markov Model, we then demonstrate that this event-specific gaze behavior is reflected in the multi-voxel MRI pattern of the eyes, and that these eye voxel patterns generalize across viewing and recall, supporting our second prediction. 5) Finally, we link the behavioral and neural domain directly by relating the eye voxel pattern to brain activity using deep learning-based gaze predictions. In line with our third prediction, we found widespread gaze-dependent modulations of brain activity that overlapped substantially between viewing and recall. We conclude by discussing these results in the context of existing theories of gaze reinstatement and neural reactivation, and outline a parsimonious theory for their relationship.

### 1) Study design and empirical measures

The present study features two datasets with independent participants, one acquired inside an MRI scanner (Dataset 1, the original “Sherlock Movie Watching Dataset”, n=16, (36)), the other one acquired on a desktop setup (Dataset 2, n=21). All participants viewed and then verbally recalled an episode of the BBC show “Sherlock” (48 minutes of the first episode: “A Study in Pink”, split into 2 acquisition runs, Fig.1A). Participants were instructed to describe the movie for as long as they wished, in as much detail as possible, while maintaining the chronological order of events. In both datasets, eye-tracking data were collected during movie viewing, as well as audio recordings to capture subsequent spoken recall. In addition, Dataset 1 included the fMRI data for which robust neural reactivations were already reported earlier (e.g., (36, 37)). For more details, see methods and data overview (Fig.1B).

**Figure 1:**
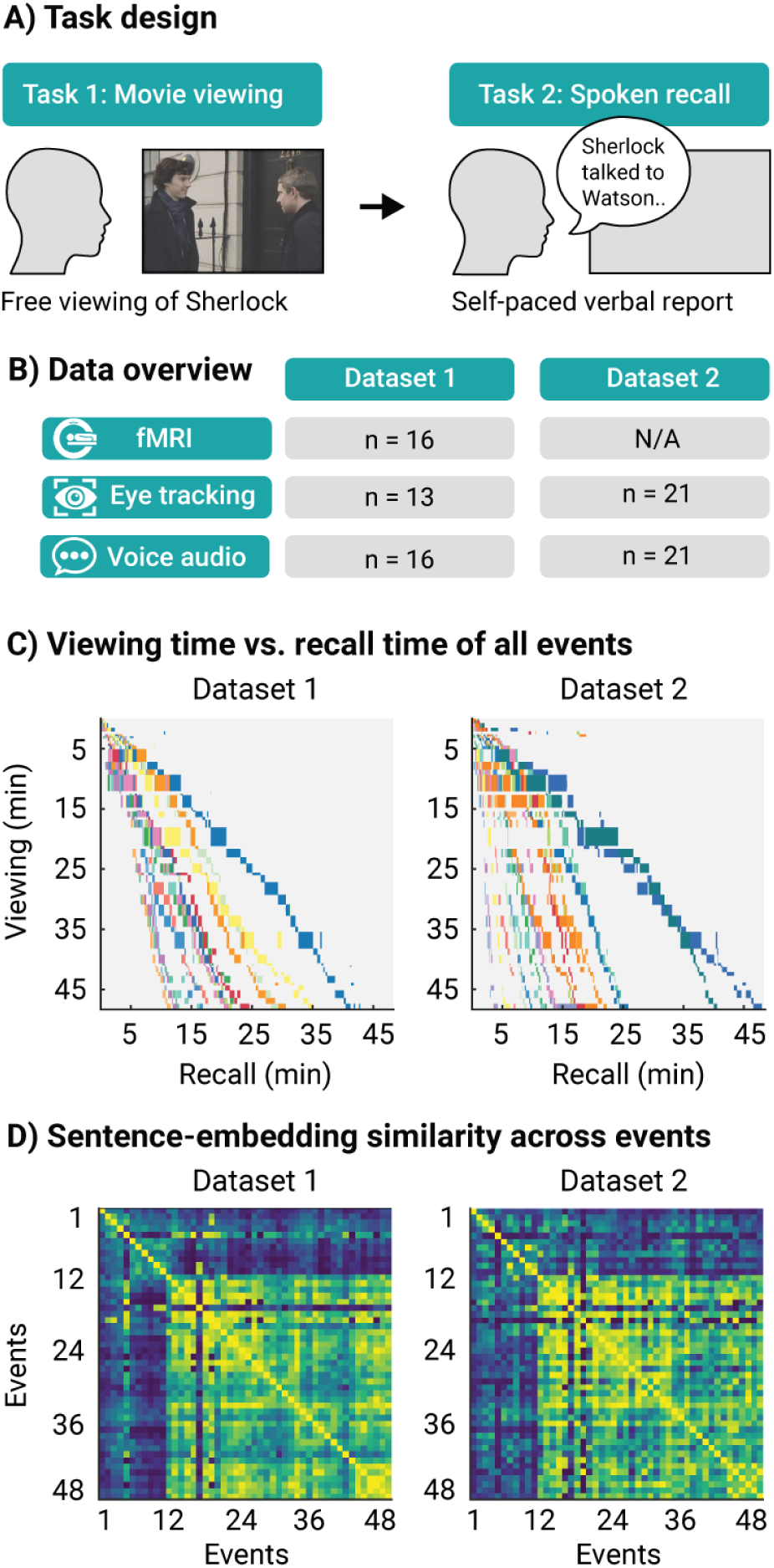
Study design and memory performance. A) Task design. Participants first viewed 48 minutes of the BBC show “Sherlock” (Task 1) and then recalled it verbally (Task 2). The movie clip was segmented into 48 narrative events by an independent viewer (36). B) Dataset overview. Eyetracking data and voice recordings were acquired for two datasets in independent participants. In addition, Dataset 1 included simultaneously acquired fMRI data. Table shows participant counts for the respective data types. C) Recall quantification 1: Summary of the order and duration of the events in both tasks. Each box represents an event and is scaled according to the duration in the respective task. Participants were color-coded within each dataset. D) Recall quantification 2: Language-model results of the spoken recall data expressed as event-by-event sentence-embedding similarity. Matrices show the ranks of Pearson correlations between sentence embeddings estimated for each event within each participant, averaged across them. Ranking was performed after computing the correlations to visually highlight similarities between matrices (blue to yellow colors show low to high ranks). Spoken recall was highly similar between the datasets.

### 2) Spoken recall follows the event structure of the movie

To ensure that the movie was recalled accurately and in enough detail for event-specific analyses, we analyzed participants’ spoken recall using a language model. To do so, we transcribed the audio files to text, followed by manual segmentation into 48 narrative events. These events were previously defined by an independent coder and reflected key, separable elements of the narrative (see (36) for details). For each narrative event recalled by a participant, we generated its embeddings using the sBERT language model (50). These embeddings are numerical representations of sentences in a high-dimensional space, allowing for the comparison of sentence meanings. The embeddings were then compared with all other recalled events using Pearson correlation.

We found that the order and duration of recalled events closely matched that of the actual events in the movie in both datasets (Fig. 1C), even though the recall tended to be compressed in time (similar to results by (51)). In addition, we observed that the semantic structure of the movie (i.e., the pair-wise similarity in sentence embeddings between all events) replicated remarkably well across datasets, and was generally consistent across participants (similar to results by (45)). By further comparing these recall results to those obtained for a “ground-truth” description of the movie created by an independent participant (Fig. S1, (36)), we found that the events were indeed recalled with high accuracy not only in terms of their relative order in the movie, but also in terms of their semantic content.

### 3) Gaze behavior is event-specific and robust across participants

Having established that our data were well suited to study event-specific processes such as those posited to underlie gaze reinstatement and neural reactivation, we next focused on the eye tracking in both datasets. These data were collected using infra-red camera-based eye trackers and denoised prior to analysis (i.e., outlier removal, detrending, and smoothing, see methods). Note that in Dataset 1, the eye-tracking data were acquired together with the fMRI data during scanning. For each event, we then computed the average saccade rate, amplitude, and duration during movie viewing, as well as a heatmap of gaze positions averaged across all frames of the respective event. These heatmaps were then compared across events using Pearson correlation to obtain similarity matrices analogous in structure to those obtained for spoken recall using the language model (Fig. 1D).

If gaze reinstatement and neural reactivation are related, gaze patterns in our data should be consistent across participants and specific to narrative events, like neural activity ((36, 52), *prediction 1*). Visualizing the eye tracking time series indeed revealed a high consistency across participants, both across in-scanner and out-of-scanner settings (Fig. 2A). We further confirmed this consistency by correlating the time series across participants within each dataset, finding robust rank correlations throughout (Dataset 1: rho = 0.53, Dataset 2: rho = 0.63, average across pairwise comparisons between participants). In addition, we observed substantial variability in saccade parameters across narrative events, and that this variability was shared across the two datasets (Fig. 2B). In other words, an event with a high saccade rate in one dataset also exhibited a high saccade rate in the other dataset. Finally, the pair-wise correlations in heatmaps across events revealed a similar pattern of results for both datasets (Fig. 2C).

**Figure 2:**
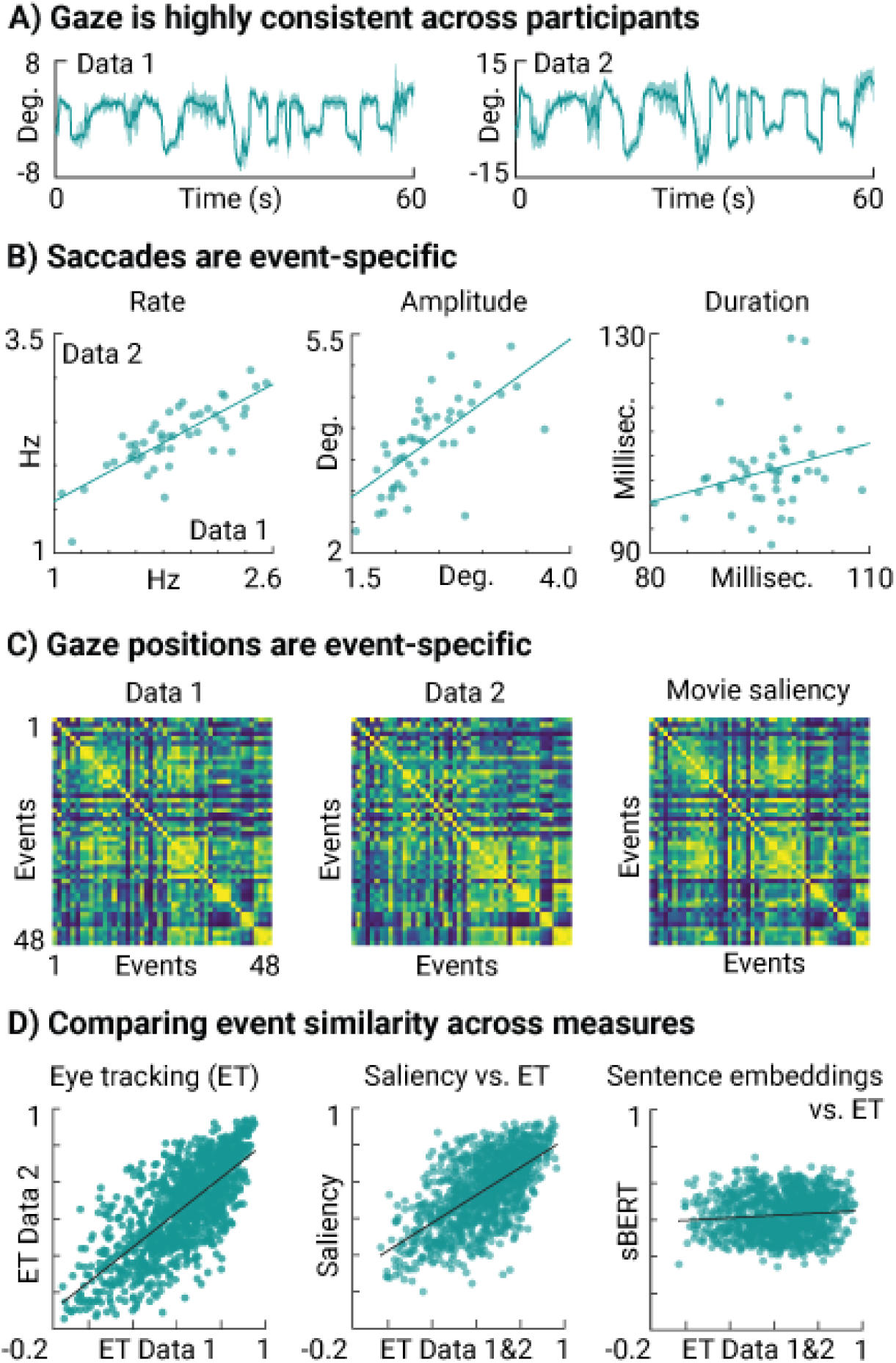
Viewing behavior is event-specific and consistent across participants. A) Group-level gaze example. Horizontal gaze position averaged across participants (green line) with standard deviation (shaded area) for Dataset 1 (left) and Dataset 2 (right). Gaze behavior was highly consistent across participants and datasets. B) Saccade parameters: Scatter plots show saccade rate, amplitude, and duration for each narrative event averaged across participants (green dots) for both datasets. Regression lines added. C) Eventby-event gaze-map similarity for Dataset 1 (left) and Dataset 2 (middle), as well as for model-derived saliency (53) of the movie stimulus (right). Matrices show the ranks of Pearson correlations between gaze maps obtained for each event. Ranking was performed after computing the correlations to normalize matrix range (blue to yellow colors show low to high ranks). Saccades and gaze positions were event specific and robust across datasets. D) Event-specific gaze patterns are highly reliable and explained by frame-wise saliency, not sentence embeddings. Scatter plots show the relationship between the lower diagonals of the event-by-event similarity matrices in Fig. 2C and Fig. 1D: Eye-tracking Dataset 1 vs. 2 (left, r = 0.73), movie saliency vs. averaged eye tracking (middle, r = 0.64), averaged sentence embeddings vs. averaged eye tracking (right, r = 0.1). Regression line added.

While these results demonstrate that gaze patterns during movie viewing are event-specific and robust across participants, confirming *prediction 1*, the pattern of results seemed to differ from the semantic similarity estimated for the spoken recall (e.g., based on the visual comparison of Fig. 1D and Fig. 2C). This is noteworthy since the events were originally defined based on narrative elements of the movie. To better understand the difference between gaze and spoken recall, we therefore modeled the saliency of each movie frame using a gaze-prediction model (DeepGaze IIE, (53)), and then compared the average saliency across events. A strikingly similar pattern emerged as the one observed for the eye-tracking data (Fig. 2C), implying that the event specificity of gaze patterns is largely explained by the visual content of the movie, not its narrative content (see Fig. 2D for direct comparison).

Importantly, all eye-tracking analyses presented so far were obtained for movie viewing. For recall, no eye-tracking data were collected inside the MRI scanner, and acquiring them on a desktop setup proved difficult (e.g. participants tended to look away from the screen and outside the range of the camera, see “Discussion” section). For all subsequent analyses, we therefore turned from camera-based to magnetic resonance-based eye tracking. This approach inherently builds on the fact that the eyeball orientation and movements strongly affect the multi-voxel pattern of the eyes measured with MRI (49). Therefore, the eyeball MRI-signal, or MReye signal for short, allows inferring gaze related variables even in existing fMRI datasets such as ours (Fig. S2).

### 4) Event-specific gaze patterns are recapitulated during recall

Using the MReye signal, we next tested whether gaze patterns observed during movie viewing were indeed recapitulated during recall. To do so, we employed a Hidden Markov model (HMM) approach previously shown to uncover event-specific brain activity and neural reactivation in fMRI data (37, 54). Critically, here, we used this approach to model the multi-voxel pattern of the eyes rather than of the brain (Fig. 3A), in order to test for evidence of concurrent gaze reinstatement. Based on our previous observations (Fig. 1, Fig. 2, Fig. S2), we reasoned that the HMM should in principle be able to learn the event structure of the movie from the eyes as well.

**Figure 3:**
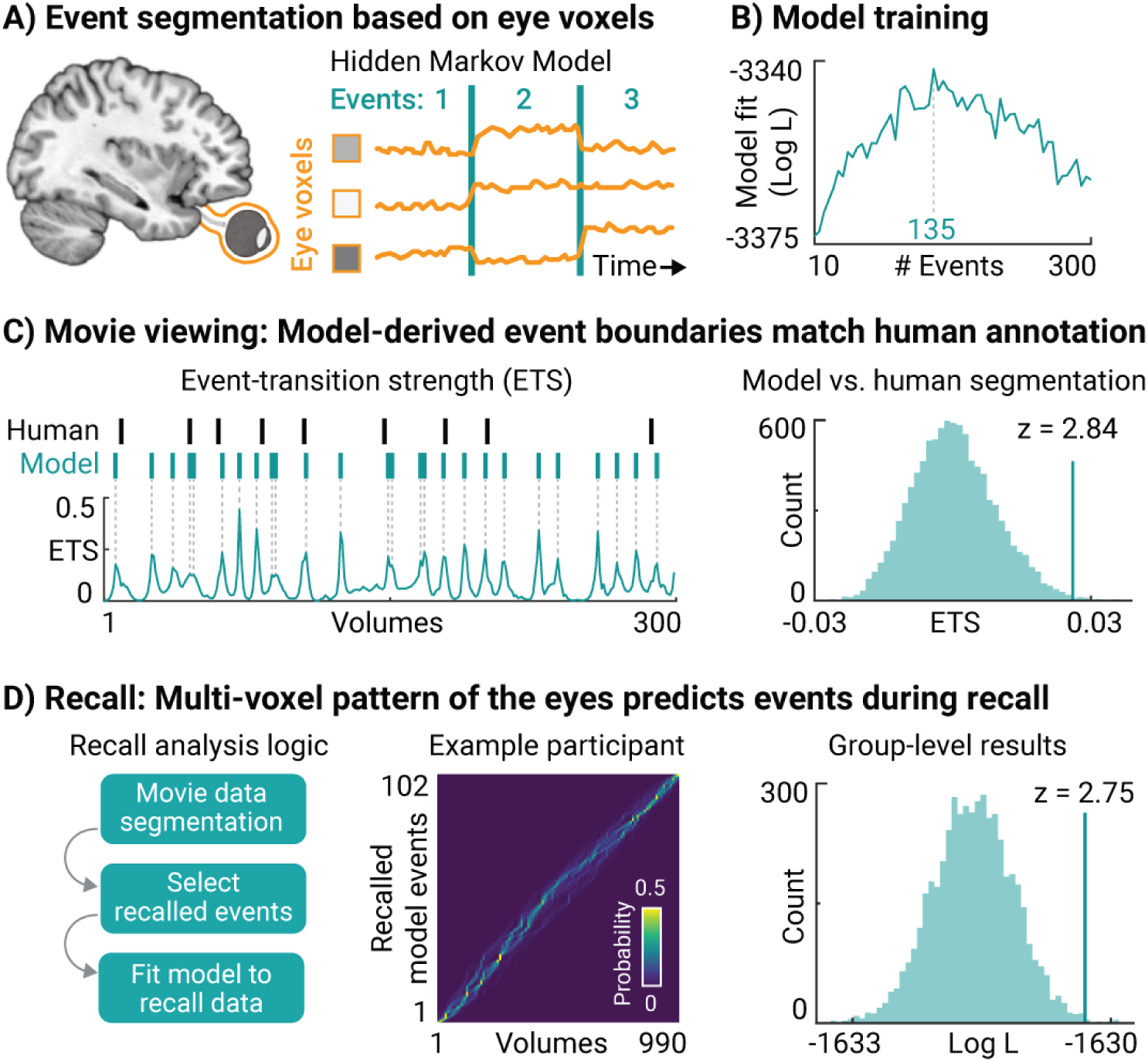
Eye voxel-based event segmentation reveals evidence for gaze reinstatement. A) Hidden Markov Model (HMM). We trained a HMM to segment the eye-voxel time series acquired during movie viewing into discrete events defined by temporally stable multi-voxel patterns. Once trained on movie viewing, we tested the model on data acquired during recall. B) Model training. We fit the HMM to the data of half of the participants, and tested it on the other half, in order to obtain a cross-validated, log-scaled model fit score (Log L). Repeating this procedure for a range of specified number of events (10–300) revealed a maximal model fit for 135 events. We therefore fit the final HMM with 135 events to the full participant pool using the movie viewing data. C) Model vs. human event segmentation. For each of the 48 human-defined event boundaries, we computed the model’s event-transition strength during movie viewing. The average event-transition strength was higher at human-defined event boundaries compared to a shuffled distribution, which was obtained by permuting the order of events while keeping their durations constant (n=10000 shuffles). D) Recall analyses. None of the participants recalled all events. For analyzing recall, we therefore created participant-specific HMM’s that searched for events that were actually recalled by the respective participant. Fitting these individualized HMM’s with the correct event order resulted in a higher model fit compared to shuffled event orders (n=5000 shuffles). These results suggests that event-specific eye-voxel patterns observed during movie viewing were recapitulated (at least partially and in the correct order) during recall.

To test this idea, we extracted the eye voxels using an automated pipeline (49), followed by denoising of the voxel time series through nuisance regression of the head-motion estimates, linear detrending, and z-scoring (see methods). We then trained the HMM by fitting it repeatedly to the movie-viewing data using a variable number of model events (10–300), similar to earlier reports (55), finding that 135 events led to a peak cross-validation performance across two participant sub-pools. We then re-trained the model on the full participant pool using these 135 events, obtaining a highly accurate model capable of recapitulating even human-defined event boundaries in our data (z = 2.84, p = 0.005, n=10000 shuffles, Fig. 3C). This successful model training demonstrates that it was indeed possible to segment the movie into meaningful events based on the MReye signal using the same techniques employed to study event-segmentation processes in the brain (37, 54).

Importantly, not every participant recalled every event (Fig. 1C). When testing whether gaze patterns generalize across viewing and recall, we therefore ensured that the trained HMM only searched for events that were actually recalled by the respective participant. To this aim, we created participantspecific copies of the trained HMM, and then removed the events that the respective participant did not recall from their individualized model (Fig. 3D, left panel). The participant-specific HMMs were then fit to the recall data, predicting which event the participant was recalling at every moment in time (Fig. 3D, middle panel). In line with the idea that gaze patterns were at least partially reinstated, we found that model performance was higher for the correct order of events compared to shuffied orders (z = 2.75, p = 0.003, n=5000 shuffles), similar to results obtained for brain activity in the same data (37). Note that similar results were observed even without limiting the analyses to the events that were recalled (z = 2.04, p = 0.021), or when the model was specifically trained on finding 48 events to match the human annotation (z = 2.03, p = 0.021). These control analyses suggest that the generalizable patterns we found in the MReye signal are robust across model-training schemes.

### 5) Gaze-dependent brain activity overlaps between viewing and recall

The results presented so far provide evidence in support of the first two predictions: Like neural activity (36, 37, 52), gaze patterns were robust across participants (Fig. 2A), specific to narrative events (Fig. 2BC), and generalizable across viewing and recall (Fig. 3). To test our final prediction that the behavioral and neural domain are linked, we additionally related eye-voxel derived gaze estimates to the fMRI activity recorded in the brain.

Our approach centered on decoding gaze-position estimates from the MReye signal using a deep learning-based gaze prediction framework (DeepMReye, (49), Fig. S2), and then converting these position estimates to eye-movement estimates (i.e., the vector length between subsequent positions). Moreover, the same eye-movement index was computed for camera-based eye tracking for later comparison (Fig. 4, Fig. S3). This approach resulted in a gaze predictor modeling eyemovement amplitude, which was then convolved with the hemodynamic response function, normalized, and fit to the time series of each brain voxel using mass-univariate general linear models (incl. nuisance regression of head-motion parameters).

**Figure 4:**
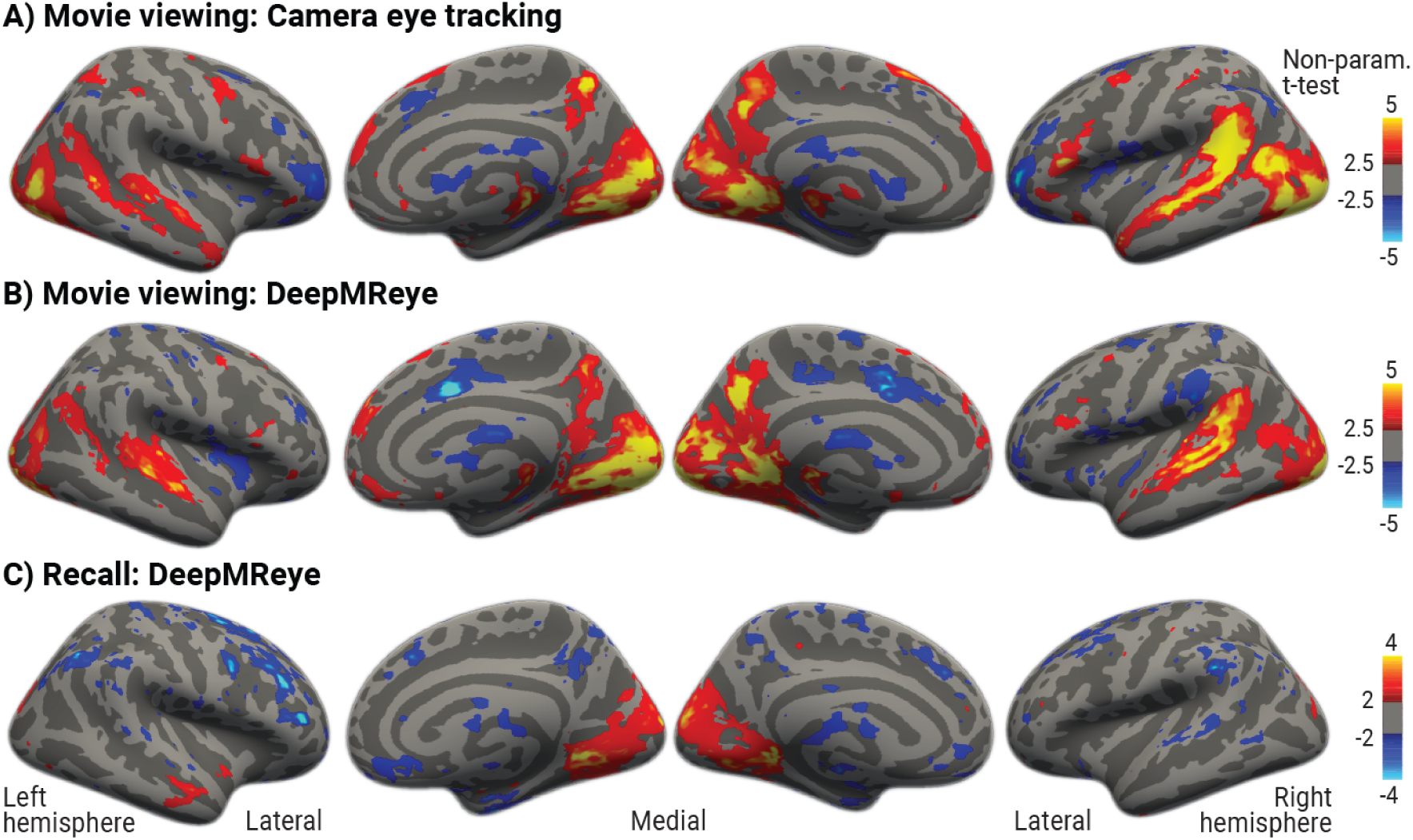
Wide-spread gaze-dependent brain activity during movie viewing and recall. All figures show voxelwise general linear model results estimated for gaze predictors modeling eye-movement amplitude (i.e., vector length between gaze positions measured or decoded for subsequent functional volumes). Statistical maps show group-level results of non-parametric, one-tailed, one-sample t-tests performed against zero overlaid on FreeSurfer’s FSaverage surface. Results are shown at liberal t-thresholds to show the spatial distribution of effects underlying the pattern in Fig. 5. A) Results obtained for movie viewing using camera-based eyetracking data (n=13). BC) Results obtained for movie viewing (B) and recall (C) using MR-based eye-tracking data (n=16) decoded using DeepMReye (49). Gaze-position changes correlate with brain activity in both tasks.

We found that the gaze predictor indeed correlated with brain activity in a wide-spread network of regions, including much of the occipital and medial parietal lobe, as well as superior and medial temporal cortices and the prefrontal cortex (Fig. 4, for volumetric, unthresholded visualization see Fig. S3). During movie viewing, camera-based and eye-voxel derived measures led to a highly similar pattern of results (Fig. 4A vs. B). However, during recall, only the eye-voxel derived gaze predictors were available. Notably, these predictors revealed evidence for gaze-dependent activity even during recall in the absence of the movie stimulus (Fig. 4C).

Importantly, if gaze reinstatement and neural reactivation are related, we expect gaze-dependent brain activity to overlap between viewing and recall. We tested this idea by computing a searchlightbased local similarity score that compared the (unthresholded and volumetric) statistical grouplevel maps obtained for the two tasks (Fig. S3B vs. C, see methods). In short, we centered a sphere with a radius of 3 voxels on each voxel to select local multi-voxel patterns that were then compared across tasks using Pearson correlation. The resulting similarity score was then assigned to each center voxel.

Using this local-similarity metric, we found evidence for strong and wide-spread overlap in gaze-dependent brain activity across viewing and recall (Fig. 5, Fig S4), confirming *prediction 3*. Moreover, a striking posterior-to-anterior sign inversion was observed on the cortical surface (Fig. 5). Specifically, gaze-dependent activity was highly similar between viewing and recall in occipital and parahippocampal cortices, whereas it was highly dissimilar in anterior parietal cortices as well as the prefrontal cortex (Fig. 5).

**Figure 5:**
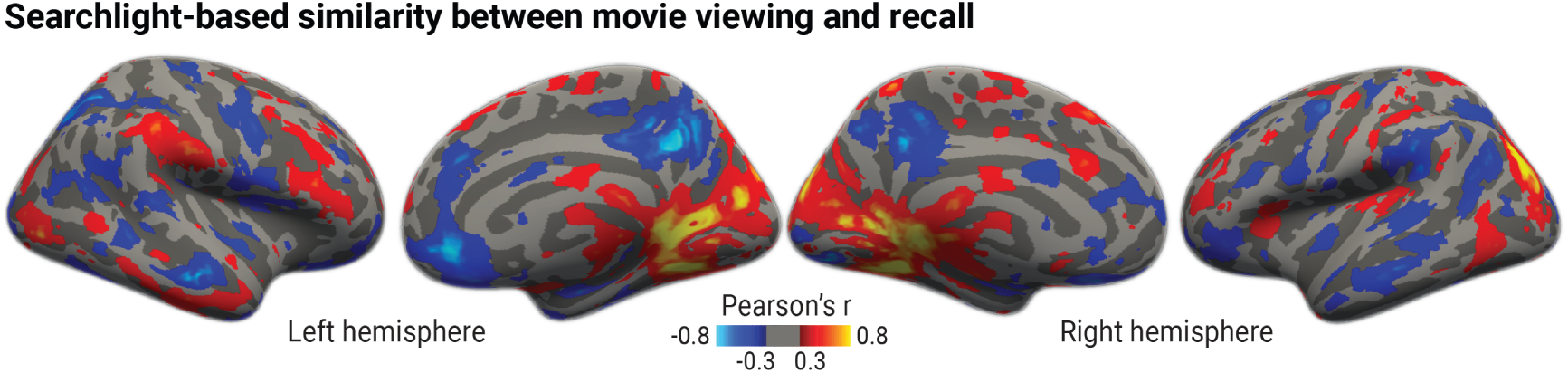
Searchlight-based similarity between viewing and recall. We centered a Xmm sphere on each voxel to select a local multi-voxel pattern, which was then compared across viewing and recall using Spearman correlation. This procedure was repeated for all voxels of the brain in volumetric space using unthresholded versions of the maps shown in Fig. 4B and C. The resulting correlation was assigned to the center voxel, and the final map was thresholded at r=±0.3 before inflating it to the FSaverage surface.

Note that, for the sake of interpretability, our DeepMReye-derived gaze predictor focused on the amplitude of (putative) eye movements, defined as the change in average gaze position across volumes. However, eye movements may affect the MReye signal even if the average gaze position remains unchanged (49). To infer dynamics in gaze behavior more generally, we therefore developed an additional, unsupervised approach based on time-varying multi-voxel pattern analysis of the eyes (Fig. S4A). Instead of training a model as for the HMM approach (Fig. 3) and DeepMReye (Fig. 4), here, a gaze predictor was created simply by computing the Pearson correlation between denoised eye-voxel patterns of subsequent volumes. This gaze predictor was then related to brain activity as described before (incl. general linear model fitting, Fig. 4, and searchlight analyses, Fig. 5, see methods). Using this approach, we further confirmed the presence of gaze-dependent activity in our data (Fig. S4B-D), finding even stronger and more wide-spread overlap in gaze-dependent activity than with our decoding approach.

## Discussion

The present study tested the hypothesis that neural reactivation and gaze reinstatement are linked through a common process underlying the reinstatement of past experiences during memory retrieval. We probed multiple key predictions arising from this hypothesis based on the viewing and recall of a narrative movie – by complementing the widely used “Sherlock Movie Watching Dataset” (36) with crucial measures of gaze behavior. In support of our predictions, we found that gaze patterns during movie viewing were event-specific and consistent across participants, thus adhering to the same principles as neural activity (36, 52). Moreover, gaze patterns and brain activity generalized across viewing and recall simultaneously within our data, with gaze-dependent activity overlapping substantially between the two tasks. Collectively, these results provide evidence that gaze reinstatement and neural reactivation are indeed deeply related phenomena, and that recall of narratives engages the same mechanisms that direct our eyes during natural vision. In addition to these conceptual advances, the present work establishes multiple new techniques and resources to leverage existing data for studying gaze-dependent activity with fMRI (see “Code availability” and “Data availability”).

### Active vision and memory retrieval are mutually constrained

Most theories of neural reactivation share the core idea that re-expressing neural patterns associated with a specific experience reinstates that experience, or at least aspects of it (see e.g., (17, 18)). This idea also extends to imagination and dreaming, likely involving the flexible recombination of patterns associated with different past experiences. Substantial empirical support for these ideas comes from studies demonstrating that the neural substrates engaged in viewing and recall overlap substantially, including reports of event-specific neural reactivation during recall of continuous narratives (36, 37, 39, 45–48).

Our results are in line with these reports and theories, while further suggesting that active vision offers a useful and complementary perspective for understanding retrieval. The conceptual starting point of the present study was the acknowledgement that the functional organization of neural circuits constrains their engagement in any and all tasks, and that many regions, including those commonly associated with memory (5, 7), are shaped by their engagement during natural vision. This engagement naturally involves gaze behavior as all visual impressions depend on it (1–3), as evidenced for example by the fact that visual field defects cause adaptive changes in eye movements (56). Moreover, we should only expect to find evidence for reinstatement of visual details that were sampled, not those that were ignored, again demonstrating that considering gaze behavior can greatly inform our understanding of retrieval. Many viewing-related principles should therefore generalize to recall and other ‘non-visual’ tasks, both on the level of neural activity and behavior, as long as task demands are shared (16).

During natural vision, our eyes move multiple times per second (1–3), each time fixating on a different aspect of the environment. Given the premise that this dynamic shapes the functional organization of neural circuits, and that the same circuits support episodic simulation (17, 18), it seems plausible that self-generated experiences follow a similar dynamic as well. For example, when experiencing our kitchen visually during recall, we may not retrieve a holistic impression of it all at once. Instead, we may retrieve individual aspects in quick succession, similar to fixations, dynamically constructing an experience that resembles viewing. This sequential retrieval would then be observable in a broad spectrum of neural sequences (for review, see e.g., (57)), even for continues experiences such as those probed here.

### Why are gaze patterns recapitulated during retrieval?

While activity patterns during recall may be constrained by those expressed during viewing, this constraint alone does not explain why gaze patterns themselves are re-expressed during recall. Active vision may again provide a useful perspective on this question: given that the underlying circuits are adapted to support vision in the context of frequent movements, it is likely that similar “sampling” mechanisms are at play when these circuits are engaged in other tasks, such as recall.

During natural vision, fixations are linked through eye movements, which need to be planned and executed based on the current retinal input, with concomitant changes in sensory experience. If recall engages similar mechanisms, reactivated activity patterns may likewise naturally trigger eye movements, indicating shifts from one retrieved item to another. Our results support this idea, for example by showing that gaze and neural patterns adhere to many of the same principles (e.g., event specificity, Fig. 2), and that eye-voxel patterns carried information about the movie event structure even though recall of visual details was not explicitly instructed (Fig. 3). Moreover, modeling eye movements revealed similar anterior vs. posterior distinctions in medial parietal activity (Fig. 5) as modeling scene viewing vs. recall (8), again highlighting the tight relationship between a circuit’s task engagement and gaze behavior.

While speculative, these considerations dovetail well with existing theories of gaze reinstatement. Scan-path theory, for instance, posits that the sequence of fixations and saccades is itself encoded, and later retrieved, as part of the memory (58). In contrast, rather than being part of the memory itself, an alternative theory posits a role of eye movements specifically in the process of retrieval (14), reinstating the “encoding context” (i.e., spatial and temporal relationships between recalled items independent of the exact scan path). A commonality between these theories and ours is that the activity patterns that drive eye movements are considered functionally relevant for memory.

Conceptualizing gaze reinstatement and neural reactivation as a consequence of shared constraints grounded in the functional organization of the nervous system not only provides a parsimonious explanation for both phenomena, but also explains why many other seemingly ‘non-visual’ tasks involve action-related signals and overt behaviors. For example, eye movements have been shown to reflect shifts between items in working-memory tasks (59–61), and the general dynamic of alternating between processing a perceptual event and shifting to another one, typically referred to as attention (62), has been linked extensively to neural activity in brain structures critical for remembering (for review see e.g., (63)).

### Toward naturalistic studies of memory retrieval

While earlier work on gaze reinstatement prioritized simple, static stimuli (14, 19–21, 24–26, 64?), we opted for movies and free viewing in favor of higher ecological validity (65). Notably, naturalistic studies such as ours face limitations with regards to the inferences about distinct experimental and cognitive factors. For example, eye movements have sensory consequences that are largely inextricable from the motor act itself, and they may correlate with other factors explaining the present data (e.g., surprise (66), fluctuations in engagement (46)). Such correlations may also explain why gaze behavior predicted brain activity in the superior temporal lobe, likely including auditory cortex (Fig. 4).

While future work may attempt to disentangle these factors, for example through behavioral encoding models (67, 68), we believe such efforts will be limited in their success. For example, these approaches implicitly assume that sensory, motor, and mnemonic signals should be fully separable on the level of neural activity – an assumption that might not be true for naturalistic paradigms such as ours (65, 69), and largely ignores the interconnected and dynamic nature of the brain (15, 16, 70, 71).

These considerations raise an important question: are eye movements a confound in studies on the neural underpinnings of memory (11)? From our perspective, the answer is no – depending on the inference made. Eye movements are not a confound, but should be considered as an inherent part of the mechanism under investigation. Rather than treating them as nuisance, it is therefore imperative to consider them when interpreting results, and to characterize their relationship to neural activity in any and all tasks (16). In line with this proposal, gaze patterns during image viewing predict later recognition performance (72–75), while restricting fixations impedes neural activity and recognition (44, 75). In addition, even without explicit retrieval (i.e., without report), neural activity predicts memory-dependent changes in gaze patterns during recognition (76).

In this context, it is also important to highlight that directed movies are designed to guide the viewer’s eyes through the scene, likely explaining the high consistency in participants’ gaze trajectories we and others observed ((52, 77–79), Fig, 2). This consistency in gaze patterns may also have contributed to the replicability of brain activity patterns across participants (36, 80). However, while gaze behavior and brain activity may be idiosyncratic outside the laboratory, the relationship between gaze reinstatement and neural reactivation may still be general.

### Advancing the study of gaze-dependent brain activity in fMRI

In addition to conceptual advances, the present work establishes multiple methods for studying gaze reinstatement and gaze-dependent brain activity in existing fMRI datasets (See “Code availability” and “Data availability”). For example, we show that the same techniques that uncover neural reactivation in the brain (Fig. 3, (37)) can be used to infer concurrent gaze reinstatement from the multi-voxel pattern of the eyes. Moreover, an unsupervised method for inferring gaze dynamics from the MRI signal of the eyes is proposed (Fig. S2), complementing earlier approaches (49, 81–84). Finally, we present two eye-tracking datasets that complement the widely-used “Sherlock Movie Watching Dataset” (36).

### Open questions

The present study quantified spoken recall in its full richness using language modeling, without acquiring specific vividness estimates of mental imagery (Fig. 1D). In principle, the strength of gaze reinstatement should correlate with self-reported imagery strength, or the number of details retrieved, which has already been shown for more constrained episodic simulation tasks (e.g., (33, 64, 85, 86)). An intriguing open question in this context is whether gaze reinstatement occurs in participants that do not report having visual imagery (i.e., aphantasics). Future studies could address such questions by systematically varying task demands (16), which have been suggested to explain variations in gaze reinstatement across studies and age groups (14).

### Conclusion

In conclusion, based on the viewing and recall of a narrative movie, we present evidence that gaze reinstatement and neural reactivation are deeply related phenomena. Patterns in gaze behavior and neural activity were event-specific, robust across participants, and generalized across viewing and recall. Gaze-dependent brain activity further overlapped substantially between the two tasks. These results suggest that viewing and recall share common constraints grounded in the functional organization of the nervous system, and highlight the importance of considering behavioral and neural reinstatement together in our understanding of how we remember.

### Data availability

The fMRI data (n=16) used in this work were shared by the original authors (36) and can be downloaded from openneuro.org: https://openneuro.org/datasets/ds001132. In addition, we share the two corresponding eye-tracking datasets, one acquired inside the MRI together with the fMRI data (n=13), and one acquired on a desktop setup (n=23) upon publication of the manuscript. The pre-trained weights used to initialize the DeepMReye for MR-based eye tracking, as well as the fine-tuned model weights we estimated can be downloaded here: https://osf.io/mrhk9/.

### Code availability

We will share the code underlying all key analyses written in Python or MatLab upon publication of the manuscript. Python code for semantic modeling of spoken recall using sBERT is publically available here: https://huggingface.co/sentence-transformers. All code for MR-based eye tracking using DeepMReye is publicly available in Python here: https://github.com/DeepMReye/DeepMReye. Python code for Hidden Markov modeling of eye voxels, as well as MatLab code for analyzing eye tracking data, and for analyzing gaze-dependent brain activity using fMRI is currently being preparing for release.

## Acknowledgements

We thank Alexandra C. Schmid for helpful discussions. M.N., A.G., and C.I.B. were supported by the Intramural Research Program of the National Institute of Mental Health (ZIAMH 002893) and the National Institute of Mental Health Clinical Study Protocol 93-M-1070 (NCT00001360). M.N. was further supported by a Feodor Lynen Research Fellowship by the Alexander von Humboldt Foundation. The funders had no role in study design, data collection and analysis, decision to publish or preparation of the article.

## Author contributions

M.N. conceptualized the present work, analyzed the data, and wrote the manuscript with supervision from C.I.B. Dataset 1 was provided by J.C., who further gave important advice on the project early on. A.G. acquired Dataset 2, transcribed the audio files to text, segmented them into events, and conducted key analyses with sBERT with supervision and code from J.A.L-V., F.P. and M.N. A.G. further helped M.N. to analyse the eye tracking data and supported the data and code release. Finally, H.T-S. and C.B. provided code and conducted preliminary analyses using the Hidden Markov model (HMM). All authors gave critical feedback and edited the final manuscript.

## Methods

### 1) Stimuli and experimental procedure

#### Movie-viewing task

Participants watched a 48 minute long segment of the first episode of the television show “Sherlock”. To allow for a short break, and to reduce the chance of technical problems, the clip was cut into two parts, one 23 minutes and one 25 minutes long. Participants were instructed to watch the episode in the way they would watch any other TV show at home, and they were told that they will need to describe afterwards what they had watched. Note that the original study (36) additionally presented a 30 second cartoon clip at the beginning of the two movieviewing sessions, which was excluded here. The Sherlock video clip itself featured rich auditory and visual content that followed an engaging narrative directed for a broad audience. Inside the MRI scanner (Dataset 1), the video was presented on a rear-projection screen using an LCD projector and subtended 20° horizontally and 11.25° vertically. Sound was presented using MRI compatible headphones. On the desktop setup (Dataset 2), the stimuli were presented on a VIEWPixx monitor and subtended 40.5° horizontally and 22.8° vertically, while the sound was presented using stereo closed-back headphones. Both experiments relied on Psychtoolbox in MATLAB for stimulus presentation (http://psychtoolbox.org/).

#### Recall task

After the two movie-viewing sessions, participants verbally described what they had watched while their voice was recorded. We refer to this session as the “Recall session”. They were instructed to recall as much detail as possible for at least 10 minutes. In the MRI scanner, the screen was black with a white central dot during recall (Dataset 1), whereas the screen was dark grey without central dot on the Desktop setup (Dataset 2). Participants were not instructed to, and did not, maintain fixation during recall. For more details, see (36).

### 2) Two datasets

Two independent datasets were used in this study with a total of 37 participants (see Fig. 1B for overview). All participants were healthy volunteers with normal or corrected-to-normal vision, gave written consent prior to data collection, and were compensated for their participation in the respective experiment.

#### Participants – Dataset 1

Publicly available functional magnetic resonance imaging (fMRI) and spoken recall data of 16 participants were downloaded from openneuro.org (https://openneuro.org/ datasets/ds001132). These data were released as part of an earlier report (36) and comprise a subset of originally tested 22 participants (10 assigned female at birth, 12 assigned male at birth, age range 18–26, all right-handed and native English speakers). Five participants were excluded from data release due to excessive head motion, whereas one was excluded due to missing data. In addition to fMRI and spoken recall data, concurrent in-scanner eye tracking data were collected in 13 of the remaining 16 participants, which are released with the present article (see “Data availability” statement). All participants gave informed consent in accordance with experimental procedures approved by the Princeton University Institutional Review Board.

#### Participants – Dataset 2

Eye tracking and spoken recall data were acquired on a desktop setup in 21 participants (13 assigned female at birth, 8 assigned male at birth, age range 22-59, all righthanded and 20 of them native English speakers). All participants gave informed consent prior to data acquisition in accordance with the guidelines of the National Institutes of Health (NIH) Institutional Review Board (National Institute of Mental Health Clinical Study Protocol NCT00001360, 93M-0170).

### 3) Spoken recall

#### Acquisition – Dataset 1 and 2

During MRI scanning, participants’ spoken recall was recorded using a customized MR-compatible microphone (FOMRI II; Optoacoustics Ltd., Dataset 1). On the desktop setup, spoken recall was recorded using a commercially available microphone (Blue Snowball USB Microphone, Dataset 2).

#### Transcription and event segmentation

The audiorecordings were transcribed to text and manually segmented into 48 narrative events, with event durations ranging between 11 seconds and 3 minutes. These events were previously defined by an independent coder without knowledge of the results or study design, and reflected key, separable elements of the movie based on location, time, and overall topic (see (36) for details). This procedure resulted in one text segment per event and participant, as well as associated time stamps reflecting the beginning and end of that event. Visualizing these time stamps showed that participants tended to recall the events in the right order in a time-compressed manner.

#### Semantic similarity with SBERT

In addition to visualizing the event time stamps, we quantified participants’ spoken recall using a language model inspired by prior work (45). Rather than comparing events in terms of their recall duration or order, or whether they were recalled or not (as shown in Fig. 1C), this approach allowed us to compare events in terms of their semantic content that was recalled. To this aim, we estimated sentence embeddings for each of the transcribed text segments using a pre-trained version of the language model SBERT (50). These sentence embeddings were then compared across events using Pearson correlation in order to obtain event-by-event similarity matrices (Fig. 1D). The pre-trained version of SBERT that was used was ‘all-mpnet-basev2’, because it had the highest average performance score for general purpose application of all pre-trained versions according to sBERT.net.

Note that SBERT models are limited to a maximum length of the text segment that is to be embedded (768 tokens), but no participants’ spoken recall ever exceeded this limit (Fig. 1D). However, some of the “ground truth” text segments of the individual events did exceed the limit. For that reason, we split them into subsegments, each of which matching the length of the shortest segment. We then compared all subsegments using Pearson correlation, and then averaged the correlations within events to obtain the event-by-event similarity matrix shown in Fig S1A.

### 4) Camera-based eye tracking

#### Acquisition – Dataset 1

During MRI scanning, eye tracking data were collected at 60Hz for 13 of the 16 participants using a long-range eye tracking system (iView X MRI-LR). Eye tracking failed in one participant due to technical difficulties during scanning, and no data was recorded in two participants despite running eye tracker. Eye tracking data were collected for the two movie-viewing scanning sessions, but not for the recall session. The data comprised position estimates that were based on pupiland corneal reflections, the latter of which was discarded due to high noise levels identified through visual inspection (e.g., strong non-physiological drift and erratic jumps to impossible tracking values). In addition to gaze position, the data included pupil size.

#### Acquisition – Dataset 2

Eye tracking data were collected on a desktop setup using an Eyelink 1000 pro eye tracking system. In 12 of the 21 participants, data were acquired at a temporal resolution of 500 Hz. The remaining 9 were recorded at 1k Hz and then downsampled to 500Hz before preprocessing. These data were collected for both movie viewing and recall, reflecting the position estimated based on the combined pupil-corneal reflection (Eyelink default). However, during re-call, participants tended to look away from the screen and outside the calibration range of the eye tracker, which rendered a big proportion of the recall data unusable. Therefore, the recall data were not considered further.

#### Preprocessing

The following steps were identical for both datasets. We denoised the eye tracking data by removing blinks in addition to outlier samples deviating more than 2x the mean-absolute-deviation (MAD) from the median gaze position. The remaining data were then linearly detrended, median centered and smoothed with a 100ms running-average kernel to remove signal drift and to further reduce noise.

#### Analysis

Multiple eye-tracking analyses were implemented. First, saccades were detected based on an eye-velocity threshold (6x MAD from the median velocity) and saccades shorter than 12 ms were excluded (87). We then computed the amplitude and duration of each saccade, and averaged them across all saccades belonging to the same narrative event. In addition, we computed the the total number (n), amplitude (amp), and duration (dur) of saccades for each functional volume.

To compare events in terms of where participants looked on the screen, we computed 2D histograms of gaze positions within each event. Each histogram contained 101 x 53 bins to match the dimensions of the stimulus. These histograms were normalized within each event and participant to sum to unity, smoothed with a 2D-Gaussian kernel with a size of 3 standard deviations, and then compared using pair-wise Pearson correlations. This procedure yielded participant-specific event-by-event similarity matrices, which were then averaged across participants to obtain one matrix per dataset (Fig. 2C). For visualization only, these group-level matrices were ranked (i.e., we converted the correlation values in the matrices to ranks), which normalizes the color scale and matches it across figures, in order to aid visual comparison.

Finally, to relate the eye-tracking data to brain activity, we computed a gaze predictor for later general-linear-model analysis (see methods section “Linking gaze to brain activity with general linear models”). For each functional MRI volume, we computed the average gaze position, resulting in a position time series that was then converted into an eye-movement time series by calculating the vector length between positions measured at subsequent volumes. For each of the movie-viewing scanning runs, a final gaze predictor was then obtained by padding the eye-movement time series with a NaN at the beginning. This gaze predictor was used to obtain the results shown in Fig. 4A. For recall, this gaze predictor could not be computed since no eye-tracking data were recorded.

### 5) Event-specific gaze patterns: semantic vs. visual content

#### Frame-wise saliency with DeepGaze IIE

Our language model-based analyses of the spoken recall (Fig. 1D) as well as our eye-tracking analyses (Fig. 2C) resulted in event-by-event similarity matrices that replicated across the two datasets. However, the pattern of results differed across the two data types, which surprised us given that the movie was segmented based on its narrative content (36). Therefore, to understand the event-specific gaze patterns we observed in more detail, we tested whether they could be predicted based on the visual content of the movie, rather than the sentence embeddings. We used a pre-trained version of the fixation-prediction model DeepGaze IIE (53) to compute the visual saliency of each movie frame expressed in the form of a 2D probability map. To reduce computational cost, we downsampled the movie to 5hz before passing it to the model. The resulting saliency maps were then averaged within each event, and then compared across events using pair-wise Pearson correlation. This procedure resulted in an event-by-event similarity matrix with the same format as the ones obtained for the camera-based eye-tracking data (Fig. 2C).

#### Comparing spoken recall, gaze, and saliency

To compare the event-by-event similarity matrices obtained for the spoken recall data (Fig. 1D), the eye-tracking data (Fig. 2C, left and middle panel), and the frame-wise saliency (Fig. 2C, right panel), we compared the lower diagonals of the respective matrices using Pearson correlation. Note that the diagonals themselves were excluded and that unranked data was used (i.e., raw, unranked versions of the matrices shown in Fig. 1D and Fig. 2C). The results of these comparisons are shown in Figure 2D (Left panel: Eye-tracking Dataset 1 vs. 2, middle panel: Average of eye-tracking Datasets 1 and 2 vs. Frame-wise saliency estimated using DeepGaze IIE, right panel: Average of eye-tracking Datasets 1 and 2 vs. Average of spoken recall Datasets 1 and 2).

### 6) Functional magnetic resonance imaging

#### Acquisition – Dataset 1

Dataset 1 included fMRI data that were acquired on a 3T Siemens Skyra using an echo-planar imaging sequence with following parameters: repetition time (TR) = 1500 ms, echo time (TE) = 28 ms, voxel size = 3.0×3.0×4.0mm, flip angle = 64°, field of view=192×192 mm. Anatomical images were acquired using a T1-weighted MPRAGE pulse sequence (0.89 mm3 resolution).

No fMRI data were collected for Dataset 2.

#### Preprocessing

MRI data were preprocessed using *fMRIprep 20.2.1* with default settings (88), making use of *FreeSurfer 6.0.1*, *FSL 5.0.9*, and *ANTs 2.3.3*. Structural scans were corrected for intensity non-uniformity using the *ANTs* function *N4BiasFieldCorrection*. Functional data were head-motion corrected by coregistering each image to a BOLD reference image computed by fMRIprep, yielding head-motion parameters estimated using *FSL’s mcflirt* function (i.e., the transformation matrix as well as six translation and rotation parameters). These data were then further coregistered to the preprocessed structural scan using *FreeSurfer’s bbregister* function with 6 degrees of freedom, and normalized to the Montreal Neurological Institute (MNI) standard space using the *ANTs* function *antsRegistration*. Using *SPM12*, these functional data were finally resampled to an isotropic voxel resolution of 3×3×3mm.

### 7) Probing gaze reinstatement with Hidden Markov Models

To examine whether the eye-voxel patterns carried information about the event structure of the movie in Dataset 1, and to see whether this information generalized across viewing and recall, we adapted a Hidden-Markov Model (HMM) approach implemented in the Brain Imaging Analysis Kit (BrainIAK, (89)). Note that this data-driven approach has been used successfully earlier for examining event-specific neural reactivations in the same dataset (37). Specifically, instead of using brain voxels, we tested whether an HMM trained on detecting events in the movie-viewing data was capable of identifying these events during recall based on the multi-voxel pattern of the eyes (Fig. 3A). Before model training, we denoised the time series of each voxel through nuisance regression of the confound time series estimated by fMRIprep (88), and by excluding voxels with an inter-subject correlation of 0.1 or lower following by previous reports (37).

#### Model training on movie viewing

In order to find the optimal number of events, we fit the HMM repeatedly to the movie-viewing data of half of the participants similar to prior work (55), each time testing a different number of events (range: 10-300). We then selected the model that led to the highest log-likelihood in the remaining half of the participants (Fig. 3B). Note that the log-likelihood is a measure of model performance and expresses how well a given model explains an observed sequence of data. Having established that the optimal number of events was 135 (Fig. 3B), we then fit the model again using 135 events to the data of all participants. This final HMM was then used for model testing.

#### Model test on movie viewing

To test whether the model segmented the movie into meaningful events that resembled the human annotation (see (36) for details), we examined whether the evidence for an event boundary (i.e., the event-transition strength, ETS) was higher at human-annotated event boundaries compared to shuffled event boundaries (Fig. 3C, left panel). Specifically, we extracted the model’s ETS at human-annotated event boundaries, and averaged them across events, in order to obtain one ETS score for the entire movie. We then shuffied the humanannotated event boundaries in time while keeping the event durations constant (n = 10000 shuffles), each time computing the score anew. We then converted the actually observed ETS score into a Z-score reflecting the score’s relationship to the shuffled distribution (Fig. 3C, right panel). Indeed, the actually observed ETS was at the tail end of the shuffled distribution, suggesting that the model-derived boundaries were more similar to the human-annotation than what was predicted by chance.

#### Model test on recall

Having established that the HMM uncovered meaningful events in the movie-viewing data, we next tested whether it could find evidence for reinstatement of these events in the eye-voxel patterns measured during recall (Fig. 3D, left panel). To do so, the HMM trained on the movie-viewing data was fit to the recall data, in order to predict which event was recalled at every functional volume. Importantly, while the HMM was trained to uncover all events in the movie-viewing data, participant’s did not necessarily recall all of these events. In fact, participants differed in which events they recalled (Fig. 1). Before model testing, we therefore created participant-specific versions of the trained HMM, each of which featuring only the events the respective participant recalled. These participant-specific HMMs were then fit to the recall data of each respective participant (Fig. 3D, middle panel), and the log-likelihood was computed as measure of model performance. Like before, we then expressed the model performance as a Z-score relative to a shuffled distribution, which we obtained by re-fitting each HMM repeatedly while shuffling the event order in the model (n=5000 shuffles). If gaze patterns were reinstated during recall, we expected model performance to be higher for the true order of events compared to a random order of events, which was indeed the case (Fig. 3D, right panel).

### 8) Magnetic resonance-based eye tracking

During MRI scanning in Dataset 1, camera-based eye-tracking data were recorded during movie viewing, but not during recall. Therefore, we used magnetic resonance-based eye tracking to infer participants’ gaze behavior from the MRI signal of the eyes in Dataset 1. The following approaches were implemented inspired by earlier work (49, 81–84).

#### Eye-voxel principal component analysis

To establish that the eye multi-voxel pattern carried information about gaze behavior in our data, we first implemented a principal component (PC) analysis using the movie-viewing data of each participant (limited to those with camera-based eyetracking). For each of the 13 participants, we estimated 10 PCs using all eye voxels and time points, resulting in 10 corresponding PC times courses. To test whether the PCs explain variance in the camera-based eye-tracking data, we used multiple linear regression to fit them to a range of gaze measures computed for each functional volume: The median horizontal (X) and vertical (Y) gaze position, the variance in horizontal (Xvar) and vertical (Yvar) gaze positions, as well as the saccade parameters reported above (saccade number, amplitude, duration). Functional volumes for which eye-tracking data were missing were excluded. Indeed, the PCs estimated for eye voxels explained substantial amount of variance in these gaze measures, especially gaze position (Fig S2A).

#### Decoding gaze position using DeepMReye

We decoded gaze position from the MRI signal of the eyes measured at each functional volume using a 3D convolutional neural network (DeepMReye, (49), Fig. S2). Using these gaze-position estimates, we then computed putative eye movements defined as the change in gaze position across subsequent functional volumes. By padding this eyemovement time series with a NaN at the beginning, this procedure resulted in one gaze predictor per scanning run similar to the one obtained for camera-based eye tracking (See methods section “Camera-based eye tracking”). However, unlike for camera-based eye tracking, this gaze predictor could be computed for both movie viewing and recall, with results shown in Fig. 4BC).

To achieve optimal model performance, we fine-tuned a pre-trained version of DeepMReye using the eye tracking data of Dataset 1. We initialized the model using weights that were estimated using a combination of guided fixations, smooth pursuit and free viewing data (https://osf.io/23t5v, weights: datasets_1to5.h5, (49)), and then fine-tuned these weights for 1 epoch with 5000 steps using the Euclidean error between measured and decoded gaze position as loss function. Following model parameters were used: batch size=8, learning rate = 0.000002, decay=0.03, no dropout, no noise. Data augmentations included 3D rotations (5 degrees), 3D translations (5 voxels), and scaling (factor=0.2).

By default, DeepMReye is trained on 10 gaze positions per functional volume (49). For model finetuning, we therefore created training labels by downsampling the eye-tracking data to 1.5 Hz, and then assigning each sample to its corresponding functional volume. If a volume comprised fewer than 50 percent valid samples, it was deemed unreliable and was excluded. Note that this procedure reduced noise, but also increased the number of missing samples per participant. Missing samples are expected for eye-tracking data in any case, especially when acquired inside the MRI scanner (e.g., calibration is more difficult than on a desktop setup). However, because gaze behavior was highly robust across participants in our data (Fig. 2), we instead fine-tuned DeepMReye using the group-level median gaze path (Fig. S2), not the data of each individual participant. Using the group-level median gaze path not only maximized the amount of training data available for each participant, but it also allowed us to use the MRI data of all 16 participants for model finetuning, instead of the 13 participants with camera-based eye tracking. Model performance was quantified as the Pearson correlation and Euclidean error between the camera-based group-level median gaze path and the decoded gaze path of each individual participant (Fig. S2).

#### Time-varying eye-voxel pattern analysis

In addition to the gaze-decoding approach described above, we implemented a new approach for inferring changes in gaze behavior based on the multivoxel pattern of the eyes. This approach is unsupervised (i.e., does not require model training) and is applicable to any fMRI dataset that comprises the eyes (Fig. S4A). The approach comprises three main steps.

First, we identified eye voxels using the automated eye extraction method implement in DeepMReye (49). Second, each voxel’s time series was denoised through nuisance regression of the head-motion parameters estimated during fMRI preprocessing, followed by linear detrending and z-scoring. Third, using these denoised voxel time series, we then computed the Pearson correlation between the multi-voxel patterns of subsequent volumes based the following logic. If gaze behavior was similar between two volumes, their pattern similarity should be high. If gaze behavior was dissimilar between volumes, their pattern similarity should be low. Consequently, and in line with previous analyses of these data (Fig. S2), the fluctuations in pattern similarity should reflect changes in gaze behavior over time. These resulting pattern-similarity time series (padded with a NaN at the beginning) was used as a gaze predictor in later general-linear-model analyses, whose results are shown in Fig. S4BC). Unlike for camera-based eye tracking, this gaze predictor could be computed for both movie viewing and recall (See methods section “Camera-based eye tracking”).

### 9) Linking gaze to brain activity with general linear models

All gaze predictors created using the camera-based and magnetic resonance-based eye-tracking techniques were related to brain activity in the same way using *SPM12* and Dataset 1. First, they were range normalized to vary between 0 and 1 in order to convolve them with the hemodynamic response function as implemented in *SPM12*. The resulting convolved gaze predictor was again range normalized and then mean-centered in order to model fluctuations around the mean of the voxel time series. Separate general linear models were fit for the different types of gaze predictors (i.e., separate models for predictors obtained through camera-based eye tracking, DeepMReye, and the time-varying eye-voxel pattern analysis). To reduce noise, the functional MRI data were spatially smoothed with 4mm before modeling (matching the voxel size). In addition to the main gaze predictors, the GLMs included head-motion parameters estimated during MRI preprocessing as well as a column of ones per run that modeled the mean of the time series.

After GLM fitting, we conducted group-level permutation-based t-tests probing whether the beta estimates obtained for the main gaze predictors were significantly greater than zero. These tests were conducted using the Statistical Non-Parametric Mapping toolbox (*SnPM*) within *SPM12* using the following settings: one-tailed, n = 10000 shuffles, variance smoothing of 6mm. The resulting group-level statistical maps were then projected to the FSaverage surface using *mri_vol2surf* and visualized using *Freeview* as implemented in *FreeSurfer 7.3.2*.

Finally, the group-level statistical maps obtained for movie viewing and recall were compared using a searchlight-based analysis. We extracted local multi-voxel patterns by centering a 3D sphere with a radius of 3 voxels on each voxel, and then compared these patterns across the two tasks using Pearson correlations. The resulting local similarity score was then assigned to the center voxel. These analyses were conducted in volumetric space using unthresholded data, and their result was again inflated to the FSaverage surface using *mri_vol2surf* for visualization in *Freeview*.

## Supplementary Material

**Figure S1:**
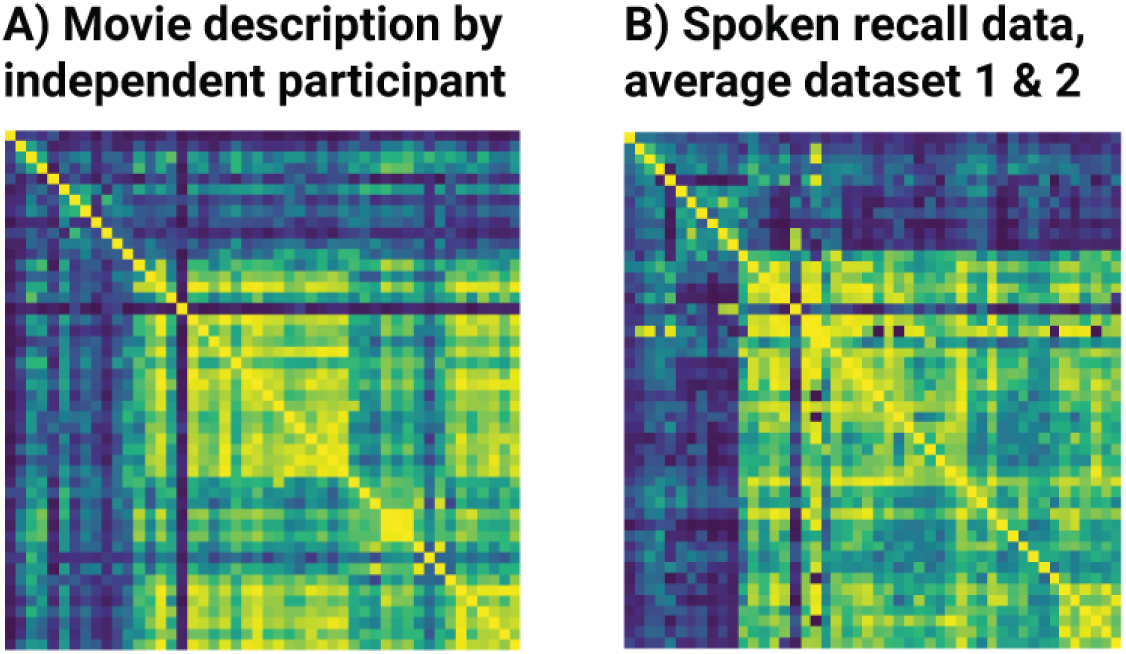
Language-model results for movie description and spoken recall. AB) Event-by-event similarity matrices depicting the ranks of Pearson correlations between sentence embeddings estimated for each event. Ranking was performed after computing the correlations to visually highlight similarities between matrices (blue to yellow colors show low to high ranks). A) Movie description created by an independent participant (36) while watching the movie. B) Spoken recall averaged across all participants of both datasets.

**Figure S2:**
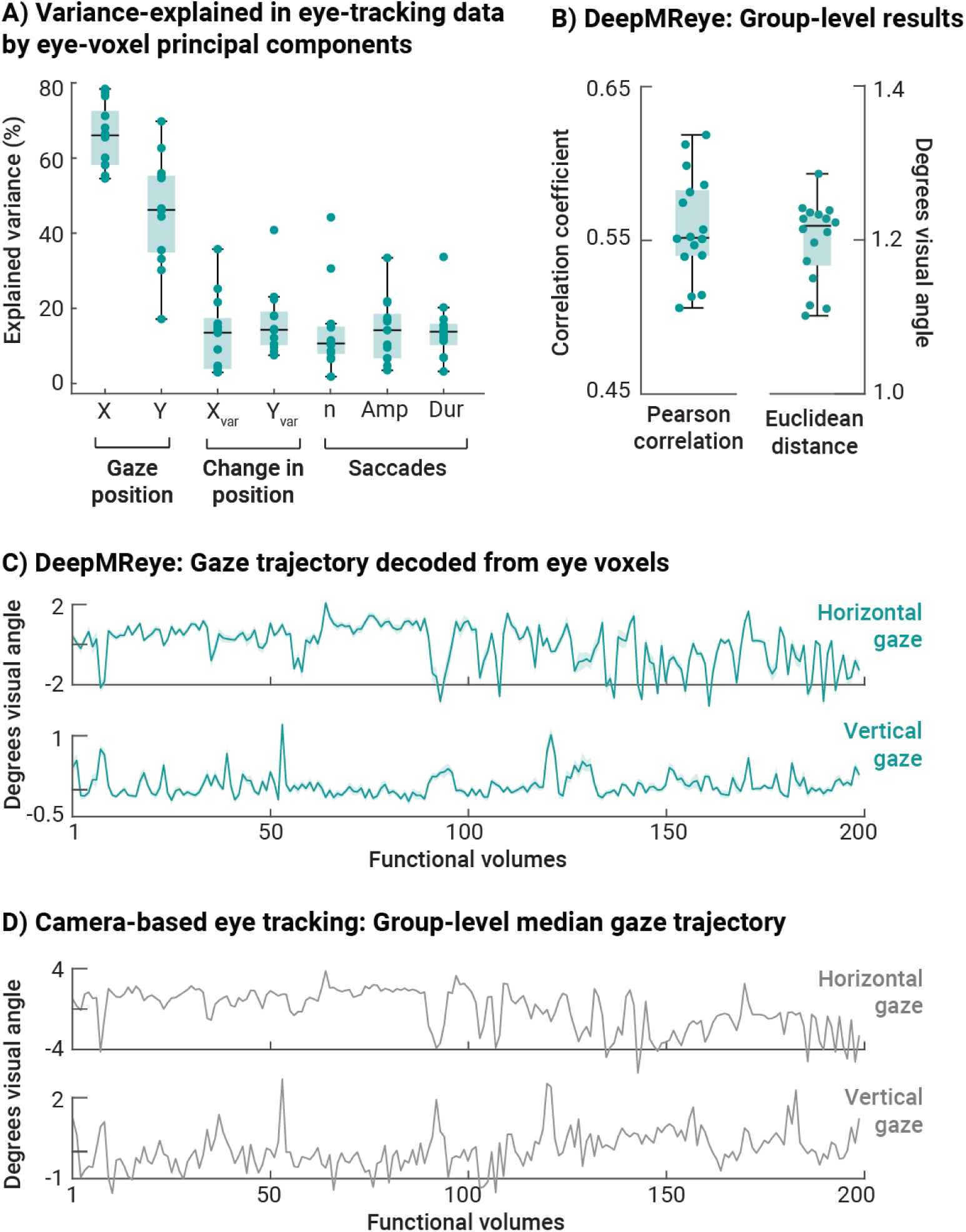
Inferring gaze behavior from multi-voxel patterns of the eyes. A) Principal component analysis. Ten principal components of the eye-voxel time series were fit to camera-based eye tracking data (i.e., horizontal (X) and vertical (Y) gaze position, variance in gaze position (Xvar, Yvar), as well as the number (n), average amplitude (Amp) and duration (Dur) of saccades. The figure shows the variance explained by the eye-voxel components in the eye-tracking measures. B) DeepMReye group-level results. Pearson correlation and Euclidean distance between the decoded gaze trajectory and the group-level median gaze trajectory (used for model training and test). AB) Whisker box plots show the median (central line), 25th and 75th percentile (box), as well as 1.5 x the interquartile range (whiskers). Single-participant data added (green dots). C) Example gaze trajectory decoded using DeepMReye shown for 200 functional volumes. We show the group-level average gaze position (dark green line) and the standard error of the mean (SEM, shaded area). Decoded gaze positions were highly similar across participants. D) Group-level median gaze position for the same 200 volumes as shown in (C). Note the similarities between (C) and (D).

**Figure S3:**
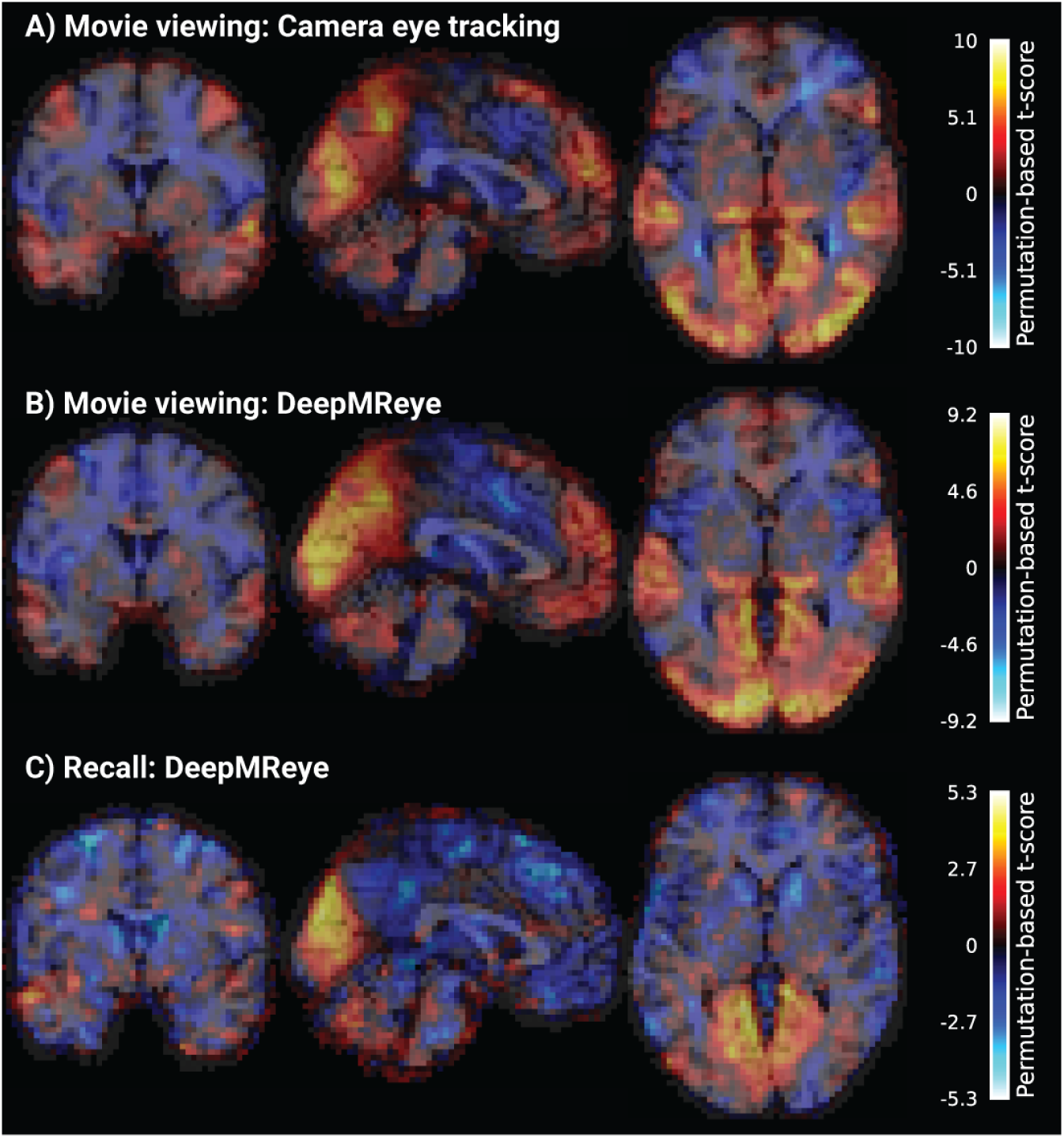
Gaze-dependent brain activity visualized without statistical thresholding and in volumentric space. All figures show voxel-wise general linear model results estimated for gaze predictors modeling eye-movement amplitude (i.e., vector length between gaze positions measured or decoded for subsequent functional volumes). Statistical maps show unthresholded group-level results of a non-parametric, one-tailed, one-sample t-test performed against zero overlaid on the volumetric Colin27 template (coordinates: X=0,Y=0,Z=0). A) Results obtained for movie viewing using camera eye tracking, BC) Results obtained for movie viewing (B) and recall (C) using MR-based eye-tracking data decoded using DeepMReye ((49)). Changes in gaze position correlate with brain activity in both tasks.

**Figure S4:**
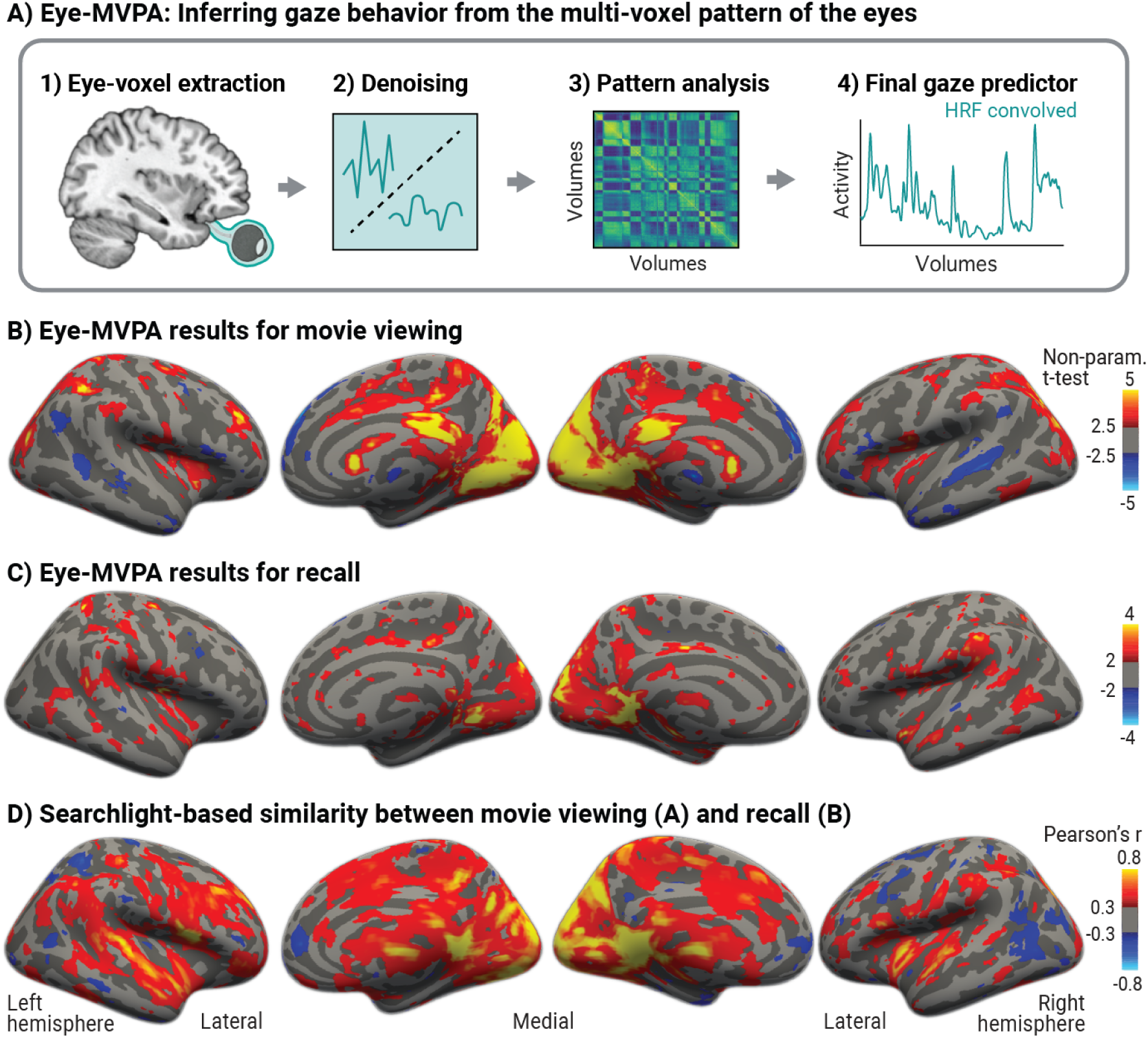
A new multi-variate pattern analysis approach to infer gaze behavior from the multi-voxel pattern of the eyes. A) Analysis logic. Eye voxels are extracted using an automated pipeline (49), followed by denoising (i.e., exclusion of out-of-eye voxels, nuisance regression of head-motion parameters, linear detrending, and z-scoring of the voxel time series). The multi-voxel pattern of subsequent functional volumes is then compared using Pearson correlation, resulting in a gaze predictor that captures the fluctuations in eye-pattern similarity over time. This gaze predictor is then range-normalized between 0 and 1, convolved with the hemodynamic response function, and then again range-normalized, before entering a voxel-wise general linear model (GLM) analysis together with nuisance regressors (e.g., head-motion parameters). BC) Results of the GLM analysis revealing wide-spread gaze-dependent brain activity during movie viewing and recall. Statistical maps show group-level results obtained for the gaze predictor using a non-parametric, one-tailed, one-sample t-test performed against zero overlaid on FreeSurfer’s FSaverage surface. Results shown for both the movie viewing (B) and recall (C) task at liberal t-thresholds to visualize the distribution of effects underlying D. D) Searchlight-based similarity between unthresholded versions of the maps shown in B and C. We centered a sphere with a radius of 3 voxels on each voxel to select the local multi-voxel pattern, which we then compared across viewing and recall using Spearman correlation. This procedure was repeated for all voxels of the brain in volumetric space. The resulting correlation map was then thresholded at rho=0.3 and rho=-0.3 before inflating it to the FSaverage surface.

